# Computational redesign of *Chlamydomonas reinhardtii* photosynthetic fructose-1,6-bisphosphatase from the Calvin-Benson-Bassham cycle improves both stability and catalytic capacities *in vitro*

**DOI:** 10.64898/2026.10.13.758731

**Authors:** Constanza Quel Piñol, Thomas Chenuel, Julien Henri

## Abstract

**Background:** Photosynthetic carbon fixation is catalyzed by 11 enzymes in the Calvin-Benson-Bassham cycle, a highly conserved metabolic pathway in the green lineage. Fructose-1,6-bisphosphatase (FBPase) contributes to the regeneration of the substrate of carboxylation. While non-photosynthetic paralogs of FBPases contribute to cytosolic neoglucogenesis in a well-documented manner, chloroplast FBPases specific regulations by phospho- and redox-posttranslational modifications, pH, cofactor binding, and homo-oligomeric exchange is only partially understood. In particular, FBPases from green microalgae are potentially target of a multiplicity of disulfide/dithiol exchanges that appear more complex than all their cytosolic and chloroplastic land-plant or cyanobacterial paralogs.

**Results:** In an attempt to decipher the molecular basis of chloroplast FBPase activities and precise redox regulations, we applied computational tools to extensively redesign its protein coding sequence, testing the simultaneous substitution of >13% of its amino acids as guided by distant homology. Resulting modified proteins in pure recombinant form were significantly more stable than the wild-type starting point, that was prone to denaturation or aggregation. In particular, redesigned FBPase favored the catalytically competent homo-tetrameric state. Thermal denaturation assays revealed that the redesign of FBPase sequence starkly increased in thermostability (+40°C), in good correlation with an augmented enzymatic activity over long periods of time.

**Conclusions:** By comparison with its wild-type counterpart, computationally redesigned FBPase allowed an extensive structure-function study that uncovered the multiplicity of redox regulations of microalgal paralog, their combined contribution to the assembly of the catalytically competent homo-tetrameric state, and will further facilitate molecular engineering of photosynthesis.

## Introduction

The metabolic phase of photosynthesis consists in carbon fixation from atmospheric CO_2_ into organic molecules, and is operated by the Calvin-Benson-Bassham cycle (CBBC, (Bassham et al., 1950),(Calvin, 1962)), a pathway highly conserved in plants and cyanobacteria. Carboxylation of ribulose-1,5-bisphosphate (RuBP) is catalyzed by ribulose bisphosphate carboxylase/oxygenase (Rubisco, (Racker, 1957),(Smillie and Fuller, 1959)) while the recycling of RuBP occurs through a reduction and the regeneration phases of the cycle. Among the 10 enzymes additionally associated with Rubisco in the CBBC, the dephosphorylation of fructose-1,6-bisphosphate (FBP) into fructose-6-phosphate (F6P) is catalyzed by fructose-1,6-bisphosphatase (EC 3.1.3.11, hereafter FBPase). FBPase is a phosphoric monoester hydrolase that, beyond photosynthesis, is also active in gluconeogenesis for the general storage of energy or for the regeneration of intermediates in central metabolism.

Molecular structure of cytosolic FBPase was first solved by X-ray crystallography (McPherson et al., 1977),(Ke et al., 1989), revealing a flat homo-tetrameric architecture and the binding sites of allosteric regulators. Later structural studies allowed for the precise mapping of the catalytic pocket, namely the identification of residues coordinating the Mg^2+^ catalytic cofactor and those forming the enzymatic pocket occupied by post-hydrolytic F6P product (Ke et al., 1990). Finally, an allosteric model of FBPase regulation was identified through structural comparison between AMP- or F6P-complexed enzyme (PDB ID: 1FBP; (Ke et al., 1991)). In this study, the authors observe that *T* (AMP-bound) to *R* (F6P-bound) allosteric transition minimally affect substrate binding pocket but actually modifies the affinity towards Mg^2+^, at a distance of 20 Å to the AMP binding pocket. The structure of an FBPase photosynthetic paralog from the chloroplast of *Spinacia oleracea* (PDB ID: 1SPI, (Villeret et al., 1995)) confirmed the structural similarity expected from >40% sequence identity supporting clear homology relationships, and shared functions (pairwise alignment RMSD=1.095 Å between 1FBP and 1SPI protomers), while also suggesting a significant evolutionary divergence over the regulation mechanisms. Indeed, the spinach chloroplast isoform lacks the structural determinants for regulatory AMP binding, and paradoxically forms an AMP-independent but nonetheless *T*-inhibited state, in a distinct mode of regulation for the photosynthetic paralogs.

Early functional studies on native FBPase extracted from spinach chloroplasts confirmed the absence of AMP-type allosteric regulation, but rather that full activity required a reductive determinant that included ferredoxin and an accessory protein factor, as well as divalent Magnesium ions (Buchanan et al., 1971). Thioredoxin was later identified as the reduction intermediate between ferredoxin and target cysteine disulfides of FBPase (Wolosiuk et al., 1979),(Nishizawa and Buchanan, 1981), and more specifically thioredoxin f was identified as the most selective in the reduction of CBBC FBPase (Yokochi et al., 2019) because of its electropositive surface around redox site, that drives TRX:FBPase docking (Lemaire et al., 2018),(Lemaire et al., 2025). In pea, the redox-targeted cysteine was identified at position 153 (Jacquot et al., 1997), with TRX-mediated disulfide reduction of C153 disulfide bridge allowing for a conformational change contributing to the formation of high-affinity Magnesium site at long-range, hence scaffolding the fully productive catalytic pocket (Chiadmi et al., 1999),(Balmer et al., 2001). This redox mechanism and primary site of redox-postranslational regulation (rPTM) was confirmed in a more distant homolog of the green lineage, the moss *Physcomitrium patens*, and further expanded rPTM mode of regulation to structurally and functionally related CBBC sedoheptulose-1,7-bisphophatase (SBPase (Gütle et al., 2016),(Le Moigne et al., 2025)). Kinetic studies of native plant FBPase also finely quantified the homo-tetrameric allosteric cooperativity, a dependence on divalent Magnesium, the activating reduction at regulatory disulfides, and ambivalent responses to metabolic regulator fructose-2,6-bisphosphate (Soulié et al., 1987),(Soulie et al., 1988),(Giudici-Orticoni et al., 1990).

A generic biological system for the CBBC is the single-chloroplast green microalga *Chlamydomonas reinhardtii*, a powerful cellular and molecular model for genomics, proteomics, and metabolomics that also holds a high biotechnology potential (Shetty et al., 2026),(Guo et al., 2026),(He et al., 2026),(Küken et al., 2018). In the work reported here, we undertook the structure-function study of *Chlamydomonas reinhardtii* FBPase (CrFBPase), in order to better understand the global regulation of CBBC at all molecular scales. Mature CrFBPase of wild-type sequence (hereafter WT-FBPase) presents 9 cysteines, some being previously demonstrated targets of S-glutathionylation, S- nitrosylation, or dithiol/disulfide exchange (Michelet et al., 2013). We resorted to extensive mutagenesis of FBPase for *in vitro* manipulation of a stable version of the protein. We describe the application of the computational tool PROSS (Goldenzweig et al., 2016) that generated a synthetic protein sequence variant with predicted improved stability (hereafter PROSS-FBPase). Redesigned FBPase indeed strikingly gained a +40°C thermostability and also proved to purify in active, stable conformation, from standard system of heterologous expression. By comparing the catalytic activity, the quaternary structures, and the redox- and pH-responses of natural (WT-) versus synthetically redesigned (PROSS-) FBPases, and informed by the resolution of a crystal structure of PROSS-FBPase at 2.3 Å, we decipher an underappreciated complexity in redox regulations of FBPase, and better illuminate how the CBBC is controlled at this prominent enzymatic step.

## Methods

### Plasmids and proteins sequences

*Chlamydomonas reinhardtii* FBPase wild-type coding sequence (Uniprot entry A8IKQ0) was amplified by PCR from cDNA and inserted in multiple cloning site of pET3c-His (ampicillin resistant, Amp^R^) between restriction sites 5’-NdeI and 3’-BamHI or in pET28a-His (kanamycin resistant, Kan^R^) between restriction sites 5’-NdeI and 3’-XhoI. Structure-function studies of WT-FBPase from these 2 plasmids are the subject of parallel investigations to be published elsewhere.

PROSS variant sequence was computed according to the standard computational pipeline and reference parameters from the online server application initially accessed in August 2024 (https://pross.weizmann.ac.il/step/pross-terms/ (Goldenzweig et al., 2016)). The optimal design8 amino acid sequence was chosen, that includes 13.62% substitutions relative to wild-type coding sequence (*vide infra*). The amino acid sequence of PROSS design8 was back-translated to nucleotide coding sequence with the codon correspondence of *Escherichia coli* high-expressing genes by EMBOSS-backtranseq from EBI online server (Madeira et al., 2024).

>PROSS-FBPase_backtranseq GCGAAAACCTCTCAGACCGAACTGATCACCCTGACCCGTTGGCTGCTGGAAGAAGAACGTAAAGGTACCATCGACGGTGAACTGGCGATCGTTATCTCTGCGATCGCGCTGGCGTGCAAAATCATCGCGTCTCTGGTTAACAAAGCGGGTATCTCTAACCTGACCGGTGTTGCGGGTAACCAGAACGTTCAGGGTGAAGACCAGAAAAAACTGGACGTTCTGGCGAACGAAGTTTTCAAAAACTGCCTGCGTTCTTCTGGTCGTGTTGGTGTTATCGCGTCTGAAGAAGAAGACGTTCCGATCTTCGTTGAAGAAACCTACTCTGGTAACTACATCGTTGTTTTCGACCCGCTGGACGGTTCTTCTAACATCGACGCGGGTATCTCTGTTGGTTCTATCTTCGGTATCTACGAACCGTCTGAAGAATGCCCGCTGGACGACATGGACGACCCGCAGAAAATGGAACAGCAGTGCATCATGAACGTTTGCCAGCCGGGTTCTAACCTGGTTGCGGCGGGTTACTGCCTGTACTCTTCTTCTACCATCATGGTTCTGACCATCGGTAACGGTGTTTTCGGTTTCACCCTGGACCCGTCTGTTGGTGAATTCGTTCTGACCCACCCGAACATCAAAATCCCGGAAACCGGTAAAATCTACTCTTTCAACGAAGGTAACTACCACCTGTGGGACGACGCGGTTAAAAAATACATGGACTCTCTGAAAGACCCGAACAAATGGGACGGTAAACCGTACTCTTCTCGTTACATCGGTTCTCTGGTTGGTGACTTCCACCGTACCCTGCTGTACGGTGGTATCTACGGTTACCCGGCGGACAAAAAAAACCCGAACGGTAAACTGCGTCTGCTGTACGAATGCGCGCCGATGGCGTTCATCGCGGAACAGGCGGGTGGTCTGGCGTCTACCGGTAAA CAGCGTGTTCTGGACGTTCAGCCGGAAGACATCCACCAGCGTGTTCCGCTGTTCATCGGTTCTAAAAAAGAAGTTGAATACCTGGAATCTTTCCTGAAAAAACAC

Nucleotide coding sequence of PROSS-FBPase was synthetized *de novo* and cloned as a synthetic open reading frame into pET28a, in fusion with a poly-histidine tag at amino-terminus (Genecust, France). Recombinant protein is fused to the tag followed by a thrombin cleavage peptide, totaling 366 amino acid residues of a theoretical molecular mass 40.2 kDa per protomer and an extinction coefficient of 38,850 mol^-1^.L.cm^-1^.

### Recombinant protein expression

Competent *Escherichia coli* BL21(DE3) were transformed with pET3-WT-FBPase or pET28-PROSS-FBPase, grown in Lysogeny broth medium under corresponding antibiotic selection, and transferred to 1L medium for inducible overexpression of protein. At DO≈0.6 isopropyl-β-D-thiogalactopyranosyl (IPTG) was added at a concentration of 0.1-0.5 mmol/L, and the cultures were kept overnight at 20°C at 140 rpm. Cells were harvested by centrifugation at 4000 rcf for 20 min, the pellet was resuspended in 20 mM Tris-HCl pH 7.9, NaCl 100 mM (buffer A), and the cells were lysed by sonication pulses of 1sec/1sec at 30 % intensity output, for 3 cycles of 1 min. Resulting lysates were cleared from insoluble material by 20 min centrifugation at 20,000 rcf, and the soluble supernatant was used for chromatographic purification of protein of interest.

### Recombinant proteins purification

First-step of affinity purification was done on gravity-flow chromatography with HIS-Select Nickel Affinity resin (Sigma). After loading of soluble fraction, 2 successive washes were done with buffer A supplemented with 10 and 20 mmol/L imidazole; elution was done with buffer A supplemented with 100 and 200 mmol/L imidazole. After quality control of expression and affinity chromatography fractions on reducing denaturing poly-acrylamide gel electrophoresis (SDS-PAGE), a second step of purification was done by size-exclusion chromatography (SEC) with a Superdex200 26/600 HiLoad on an Äkta Go FPLC system in buffer A. Elution of protein of interest was monitored by absorbance at 280 nm, while the elution fractions were collected on 96 well plates. Fractions corresponding to the main elution peaks were pooled, concentrated by ultrafiltration, titrated by Nanodrop spectrophotometry at 280 nm, and stored for later experiments. Protein apparent molecular masses were estimated from SEC elution volume, according to globular calibration standards (Bio-Rad). The corresponding homo-oligomeric states were interpreted considering multiples of a protomer mass of 40 kDa.

### Thermal denaturation assay by differential scanning ffuorimetry (DSF)

Following protocol from Boivin *et al*. (Boivin et al., 2013), WT- or PROSS-FBPase concentrated at 1 mg/mL (25 µmol/L) were mixed with SYPRO orange 2X solution in a total volume of 50 µL and distributed in 96-well qPCR plate, in triplicates. Thermal denaturation curves were followed using a CFX96 Touch Real-Time PCR Detection (Bio-Rad) by increasing temperature by incremental steps of 0.5°C every 30 sec up to 95°C, monitoring live fluorescence of SYPRO in HEX channel (λ_excitation_=515-535 nm ; λ_detection_=560-580 nm). SYPRO fluorescence increased as a consequence of protein denaturation exposing core apolar residues. The melting temperature (T_m_) was computed from the minimum of the first derivative of the fluorescence versus temperature curve using the *meltcurve* function in the *qpcR* package (Ritz and Spiess, 2008) in R.

### Enzymatic assay

Dephosphorylation of β-D-fructose-1,6-bisphosphate (FBP) into β-D-fructose-6-phosphate (F6P) was quantitatively reported through subsequent isomerization of F6P into glucose-6-phosphate (G6P) by phosphohexose isomerase (PGI, Sigma), followed by the oxidation of G6P by glucose-6-phosphate dehydrogenase (G6PDH, Sigma). Oxidation of G6P is coupled to the reduction of NADP^+^ into NADPH,H^+^ that we measured by absorbance at 340 nm on Uvikon spectrophotometer in a reaction volume of 100 µL, in a reaction mixture of 0.2 mmol/L NADP^+^, 30 mmol/L Tris-HCl pH 7.9, 16 mmol/L MgSO_4_, 0.6 mmol/L FBP, 0.1 U PGI, 0.1 U G6PDH and 1 µL of FBPase at variable tested concentrations. The linearity of the assay was validated by serial dilution of FBPase. Experiments were done in triplicates. For reduced state essay, the enzyme was incubated for 30 min in 10 mmol/L of dithiothreitol (DTT).

### Crystallization

Pure recombinant PROSS-FBPase concentrated by ultrafiltration to 5-40 mg/mL was screened for crystallization in sitting-drops of 50 nL protein supplemented with 50 nL precipitant solutions from JCSG core screens I-IV (Qiagen) and equilibrated over days by vapor diffusion at 20°C. Single monocrystals initially grew in conditions JCSG I A3 (0.05 mol/L lithium sulfate, 0.05 mol/L sodium sulfate; 0.05 mol/L Tris-HCl pH 8.5; 30% (w/v) polyethylene glycol 400), JCSG I H9 (0.1 mol/L citric acid pH 3.5; 0.8 mol/L ammonium sulfate), and JCSG IV A6 (0.1 mol/L CHES pH 9.5; 30 % polyethylene glycol 400) and later in chemically related conditions. Crystallization was optimal in 5% (w/v) PEG 6000 and 0.1 mol/L citric acid pH=4.0, yielding a large bipyramidal monocrystal.

#### X-ray diffraction and structure resolution

An individual PROSS-FBPase crystal was cryo-protected in 12.5% and 25% ethylene glycol, mounted in a nylon cryo-loop, vitrified in liquid nitrogen, and submitted to X-ray diffraction at synchrotron SOLEIL beamline Proxima-1. Collected datasets were analyzed and interpreted with XDSme (https://github.com/legrandp/xdsme) and AutoPROC (Vonrhein et al., 2024). A complete dataset in space group P 4_1_2_1_2 at a resolution of 2.29 Å was phased by molecular replacement with an AlphaFold3 model of PROSS-FBPase (Abramson et al., 2024) and 2 molecules in the asymmetric unit, and refined according to statistics reported in table 2 in Phenix 2.0-5936 crystallography software suite (Liebschner et al., 2019), including Phaser-MR, AutoBuild, Fit Loops, phenix.refine; and Coot for manual macromolecular building or corrections (Casañal et al., 2020).

## Results

*Recombinant fructose bisphosphatase from Chlamydomonas reinhardtii chloroplast* We overexpressed polyhistidine-tagged *Chlamydomonas reinhardtii* fructose-1,6-bisphosphatase (WT-FBPase) in the heterologous host *Escherichia coli* BL21(DE3), using coding sequences cloned in pET plasmids. High amount of target protein could be isolated in soluble form after resuspension of cell pellet and lysis by sonication in standard buffer (30 mmol/L Tris-HCl pH 7.9, 100 mmol/L NaCl: buffer A). However, once the His-tagged protein was purified through a first step of affinity chromatography, we repeatedly observed a progressive loss of material in the form of higher-mass molecular species that eventually sedimented as insoluble precipitate.

This intrinsic instability of recombinant WT-FBPase was quantified by differential scanning fluorimetry (DSF), with a melting temperature (T_m_) of 42.65 ±2.12°C for the untreated purified protein (Table 1). Precipitation was dependent on time, in the range of hours, and could be slowed but not stopped by the addition of chemical reducing agent (dithiothreitol, DTT) or divalent magnesium (MgSO_4_) to buffer A. Consistently with stabilization against precipitation, chemical reduction of disulfide bridges by 10 mmol/L DTT caused an increase in WT-FBPase T_m_ by 12°C, at a value of 54.68 ±0.68 °C. The addition of MgSO_4_ to purified WT-FBPase caused the duplication of thermal denaturation curve into two successive steps of T_m_ ^1^=42.62 ±0.4°C and T ^2^=71.92 ±0.4°C, accounting for the stabilization of a fraction of the protein population by Magnesium. In order to stabilize the protein in this soluble, stable and monodisperse state, we proceeded to a synthetic redesign of the protein sequence by computationally guided mutagenesis.

**Table 1.** In vitro thermal denaturation measured by differential scanning fluorimetry (DSF).

| **T<sub>m</sub> (°C)** | **WT-FBPase** | **PROSS-FBPase** |
| --- | --- | --- |
| untreated protein | 42.65 ± 2.12 °C | 80.92 ± 0.52 °C<br>84.93 ± 0.07 °C |
| + FBP 0.6 mmol/L | 43.2 ± 2.1 °C | 85.61 ± 0.45 °C |
| + MgSO<sub>4</sub> 16 mmol/L | 42.62 ± 0.4 °C<br>71.92 ± 0.4 °C | 86.62 ± 0.19 °C |
| + DTT 10 mM | 54.68 ± 0.68 °C | 75.51 ± 0.3 °C |

± values are 95% confidence intervals.

**Table 2.** Crystallographic data collection and statistics of crystallographic model.

| **PROSS-FBPase** |  |
| --- | --- |
| Resolution range | 47.91-2.289 (2.35-2.29) |
| Space group | P 41 21 2 |
| Unit cell | 115.941 115.941 118.074 90 90 90 |
| Total reflections | 72827 (5442) |
| Unique reflections | 68432 (5212) |
| Multiplicity | 1.1 (1.0) |
| Completeness (%) | 98.67 (97.14) |
| Mean I/sigma(I) | 8.64 (0.51) |
| Wilson B-factor | 61.44 |
| R-merge | 1.203 e-17 (3.526 e-17) |
| R-meas | 1.701 e-17 (4.987 e-17) |
| R-pim | 1.203 e-17 (3.526 e-17) |
| CC1/2 | 1 (1) |
| CC* | 1 (1) |
| Reflections used in refinement | 36410 (2721) |
| Reflections used for R-free | 1775 (126) |
| R-work | 0.2120 (0.3714) |
| R-free | 0.2603 (0.4296) |
| Number of non-hydrogen atoms | 4839 |
| macromolecules | 4683 |
| solvent | 156 |
| Protein residues | 606 |
| RMS(bonds) | 0.004 |
| RMS(angles) | 0.79 |
| Ramachandran favored (%) | 96.30 |
| Ramachandran allowed (%) | 3.54 |
| Ramachandran outliers (%) | 0.17 |
| Rotamer outliers (%) | 0.59 |
| Clashscore | 5.24 |
| Average B-factor | 68.09 |
| macromolecules | 68.46 |
| solvent | 56.97 |

### Computational redesign of fructose bisphosphatase for increased stability

We submitted the wild-type sequence of *Chlamydomonas reinhardtii* FBPase (Uniprot entry A8IKQ0_CHLRE, Figure 1A) to the algorithmic redesign tool PROSS (the <u>P</u>rotein <u>R</u>epair <u>O</u>ne-<u>S</u>top <u>S</u>hop) (Khersonsky et al., 2018),(Weinstein et al., 2021). Briefly, PROSS is an *in silico* method based on predicted 3D structure and on massive homologous sequence alignment that aims at optimizing protein stability to increase their potential yield in heterologous expression. The workflow is fully implemented in an online pipeline (https://pross.weizmann.ac.il) that we followed according to preset parameters and otherwise recommendation guidelines. PROSS hypothetically modifies given primary sequence to diminish computed energy difference (ΔΔ*G_cal_*_c_) between the non-mutated reference (here WT-FBPase) and additive single-point substituted mutants found as variations in large sequence alignment. Far-homology search for FBPase is actually particularly effective since this enzyme is widely represented in genomes of all kingdoms, for its universal function in core metabolism.

**Figure 1.**
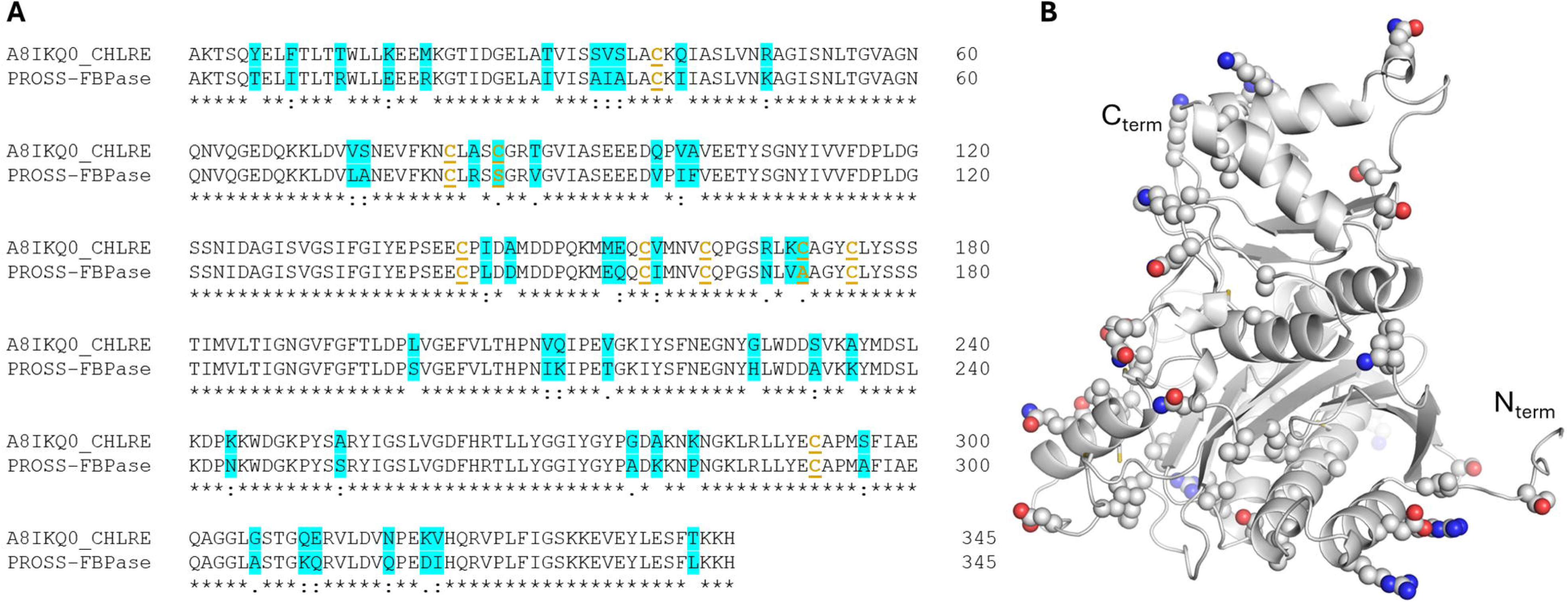
Sequence and predicted structure of FBPase from this study. A. Fructose bisphosphatase pairwise sequence alignment. A8IKQ0_CHLRE: wild-type sequence from *Chlamydomonas reinhardtii* nuclear genome. PROSS-FBPase: A8IKQ0_CHLRE-derived computational variant described in this study. 47 amino acid substitutions are highlighted in cyan. Cysteines and their substitutions are written in bold and underlined, in gold characters. B. AlphaFold3 predicted structure of PROSS-FBPase (pTM=0.93), with residues substituted shown as side chain spheres. N_term_, C_term_: amino-terminus (position 1), carboxy-terminus (position 345).

Accessed version PROSS2 has been rigorously assessed by community-wide experimental validations (Peleg et al., 2021) and proved efficient at redesigning several types of proteins including peroxidases (Barber-Zucker et al., 2022b), laccases (Barber-Zucker et al., 2022a), therapeutic glucosidase (Pokorna et al., 2023), serum albumins (Khersonsky et al., 2023), ion channel (Weinstein et al., 2024), and pathogenic adhesin (D Sa et al., 2025). To our knowledge, we report here the first use of PROSS on a chloroplastic enzyme although other approaches successfully designed *de novo* photosynthetic proteins, *e.g.* photosynthetic reaction center maquettes (Ennist et al., 2022) and artificial C_2_-symmetric, protein binders for excitonically coupled chlorophyll pairs (Ennist et al., 2024).

Following recommendation by the server, we chose design8 among PROSS-FBPase results. This variant contains 47 amino acid substitutions, corresponding to 13.6% mutations in its 345 amino acid coding sequence (Figure 1A). According to a prediction of 3D structure of PROSS-FBPase by Alphafold3 (Figure 1B), mutated residues are scattered in several positions of the protein, several being at the solvent-accessible surface and with an increase in polarity relative to WT-FBPase, other being at the protein folded core and with an increase in hydrophobicity relative to WT input. Still others amino acid modifications selected by PROSS have no obvious physicochemical advantage, *e.g.* lysine (K) to proline (P) substitution or the replacement of 2 cysteines (C) by redox-insensitive isosteric residues (respectively serine, S, and alanine, A).

### Expression and purification of highly stable redesigned fructose bisphosphatase

Notwithstanding our incomplete understanding of PROSS-suggested extensive modifications over *Chlamydomonas reinhardtii* input WT-FBPase sequence, we generated the nucleotide coding sequence for design8 PROSS-FBPase with optimal codon bias for bacterial overexpression, ordered the synthetic gene and its subsequent cloning into pET28 plasmid, and applied the protein preparation protocol described above for WT-FBPase. Redesigned PROSS-FBPase was highly overexpressed in soluble form and could be efficiently retained by its polyhistidine-tag on nickel resin, eluting in a pure state from affinity chromatography (Figure 2A, lane elution 1). Eluted protein was stable over time, with no sign of material loss nor precipitation as insoluble aggregates. Compared to its WT-counterpart, PROSS-FBPase strikingly gained 38 to 42°C in DSF-measured thermal stability, with dual behavior of the purified protein at T ^1^=80.92 ±0.52°C and T ^2^=84.93 ±0.07°C (Table 1). This result experimentally validates again the PROSS method of screening all amino acid mutations to optimize the protein computed energy with constraints inferred from homologous sequences (Goldenzweig et al., 2016). However, the duality in thermal denaturation of the redesigned protein was not expected.

**Figure 2.**
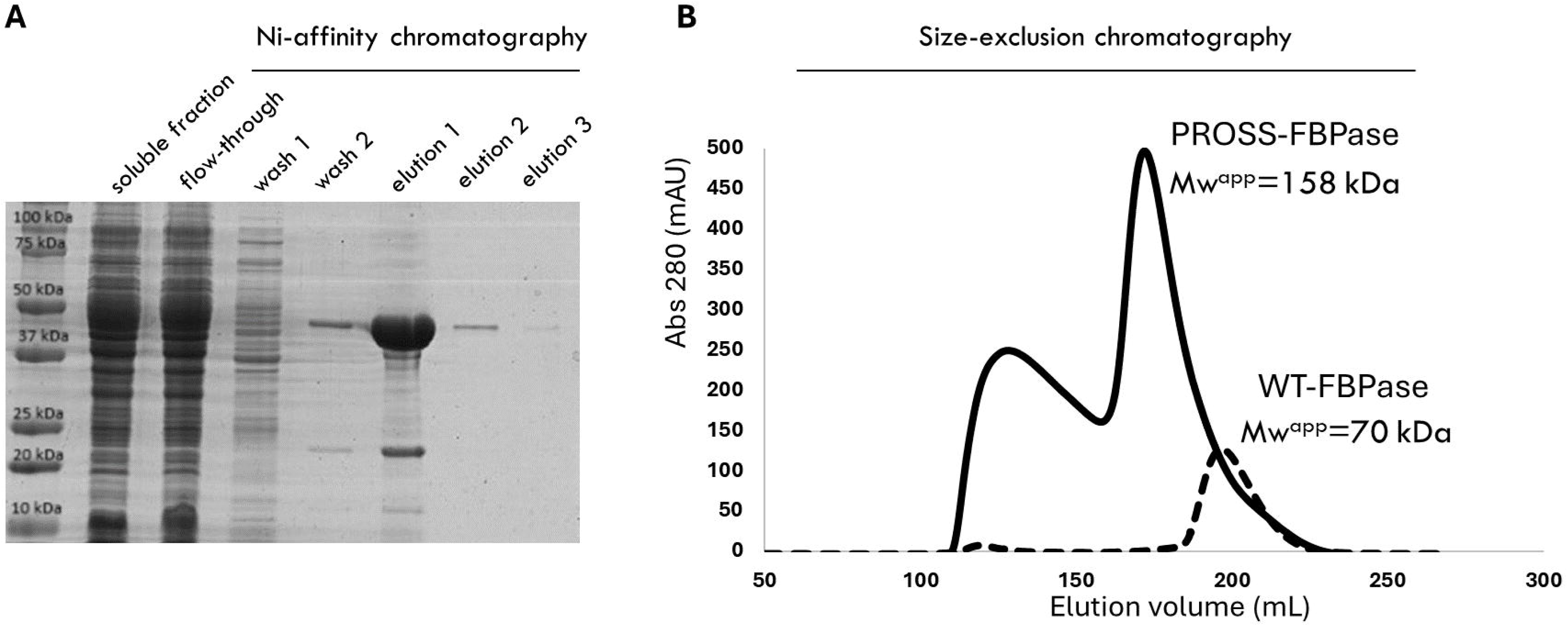
Purification of recombinant fructose bisphosphatase. A. Denaturing, reducing polyacrylamide gel electrophoresis (SDS-PAGE) of purification fractions from Nickel-affinity (Ni-NTA) chromatography. Staining by Coomassie blue. B. Superdex200 26/600 HiLoad size-exclusion chromatography of Ni-NTA elution profile. Plain line: PROSS-FBPase; dashed line: WT-FBPase. Apparent molecular masses (Mw) are computed according to globular calibration standards.

### PROSS-FBPase binds its substrate and ion cofactor

We probed the functionality of PROSS-FBPase according to its ability to bind canonical ligands, starting with the DSF measurement of its thermal stability in presence of fructose-1,6-bisphosphate (FBP). When pre-mixed with 0.6 mmol/L FBP, 25 µmol/L PROSS-FBPase thermal denaturation curve switched from dual to unimodal, with one single population of PROSS-FBPase:FBP denaturing at T_m_=85.61 ±0.45°C (Table 1). This is coherent with a mixture of the initially *apo* enzyme populating two conformations, that falls into one single population of homogenous conformation in the presence of a saturating excess of its cognate ligand. Comparatively, redox untreated and FBP-mixed WT-FBPase do not show such a switch in thermal denaturation profile after substrate addition (Table 1), suggesting that purified WT-FBPase has a low affinity towards its substrate.

Considering that PROSS-FBPase thermal stability is further increased in the presence of substrate, we consider that computational redesign maintained or even favored an active conformation of the protein. Among the 2 populations of apo protein, binding of F6P selects for a highest state of stability. Apparently saturated by its substrate, pure recombinant PROSS-FBPase forms a single population that we interpret as the Michaelis complex.

We similarly probed the effect of divalent Magnesium (FBPase catalytic cofactor) on purified PROSS-FBPase DSF denaturation curve. 16 mmol/L MgSO_4_ mixed with 25 µmol/L PROSS-FBPase also yielded a single population with T_m_=86.62 ±0.19°C, in the same range of stability as the enzyme:substrate mixture (Table 1). This confirms that multiple populations of *apo* enzyme are competent to bind Magnesium, adopting a cofactor-stabilized conformation, and thus forming a single population of higher stability. This enzyme:cofactor complex is also expected to represent the catalytically competent conformation of PROSS-FBPase that FBP also stabilized. Meanwhile, only a fraction of WT-FBPase is stabilized by Magnesium, at at lower T_m_ than PROSS-FBPase, owing to a lower affinity towards ligand and/or a more labile conformation of activated WT-FBPase.

### Enzymatic activity of PROSS-FBPase

Considering the ability of redesigned FBPase to productively interact with Magnesium cofactor and with FBP substrate, we assayed its catalytic activity *in vitro* with a spectrophotometric coupled-reporter assay (Figure 3). FBP saturation curve of the enzymatic hydrolytic activity is representative of a Michaelis-Menten hyperbolic function (Figure 3A), with apparent kinetic constants K_M_=0.23 mmol/L and k_cat_=0.1 sec^-1^. These value compare to previously reported kinetic constants of *Euglena gracilis* recombinant chloroplastic FBPases. In this other microalga, the two forms of FBPase were characterized: isozyme I with K_M_=165 µmol/L and k_cat_=0.17 sec^-1^, and isozyme II with K_M_=2,200 µmol/L and k_cat_=1.08 sec^-1^ (Ogawa et al., 2015). EgFBPase isozyme I is sensitive to redox regulation, accounting for its photosynthetic character, while isozyme II is insensitive to oxidative treatment and more probably belong to cytosolic-type of core metabolism.

**Figure 3.**
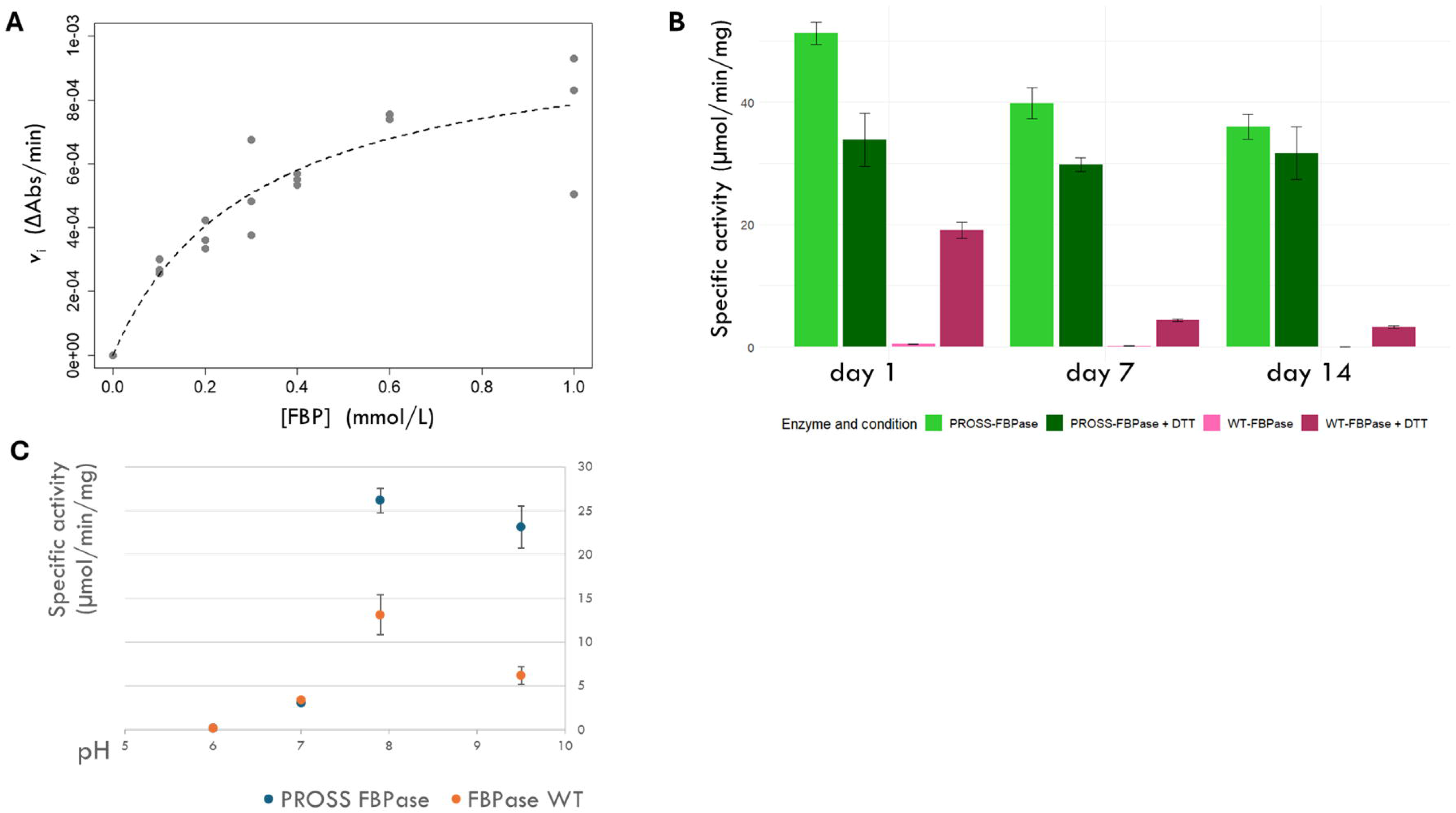
In vitro activity measured by spectrophotometric coupled assay. A. Michaelis-Menten saturation curve of fructose bisphosphatase reported activity with initial velocity (v_i_) plotted according to substrate concentration (FBP). B. Reported FBPase specific activity along time after purification, in untreated conditions or after reducing treatment by dithiothreitol (DTT). Green: PROSS-FBPase; pink: WT-FBPase. C. pH sensitivity of PROSS- and WT-FBPase.

Taken together and with DSF results, this hence validates that computationally redesigned of PROSS-FBPase maintained its ability to productively bind to Magnesium cofactor and FBP substrate, in order to perform catalysis with enzymatic constants that compare to those of a known photosynthetic FBPase from a green microalga.

### Effect of chemical reducing treatment on FBPase enzymatic activity

We tested the sensitivity of FBPase to reducing treatment with chemical agent dithiotreitol (DTT, 10 mmol/L) as an *in vitro* proxy for CBBC physiological control through the ferredoxin-thioredoxin cascade of FBPase activation. Purified WT-FBPase appeared strongly dependent on reducing treatment to reach a measurable *in vitro* activity in our reporter assay (Figure 3B), consistently with a purified protein holding inhibitory disulfide bridges. Meanwhile, DTT treatment of PROSS-FBPase caused no increase in specific activity, consistently with the purified state being devoid of inhibitory disulfide. DTT treatment of PROSS-FBPase actually moderately reduced its enzymatic activity, pointing to the presence of a disulfide bridge contributing to the stable conformation of the protein. Overall, these opposite and opposite effects of chemical reduction between the natural and redesigned FBPase accounts for a multiplicity of disulfide bridges bearing distinct effects on the active site. Recombinant WT-FBPase is purified in an auto-inhibited state due to the presence of an inhibitory disulfide bridge while recombinant PROSS-FBPase is purified in active form independent of such an inhibitory bridge and/or in the reduced state for this bridge, and instead depends on the presence of a previously invisible stabilizing disulfide bridge to reach its highest state of activity. The position of cysteine and their state of reduction will be discussed later in the context of the crystal structure of PROSS-FBPase.

Over time, PROSS-FBPase maintains a relatively high level of enzymatic potential, loosing only ∼20% of its activity every week. Interestingly, its decrease in activity caused by DTT treatment leads to a stable level activity, suggesting that the PROSS-positively contributing disulfide bond loses its stabilization effect over time. When the enzyme reaches its basal, redox-insensitive state of relatively high activity, it remains unchanged and insensitive to treatment. This illustrates a contribution of this stabilizing disulfide in the biogenesis (folding and assembly) of the active enzyme first, and later to a partial additional gain in activity maximally seen at day 1. On the contrary, DTT activation of WT-FBPase only yields a relatively low specific activity, never reaching that of PROSS-FBPase, and that appears to be irreversibly lost over time.

### Effect of reducing treatment on FBPase stability and structure

DTT treatment confirmed its complex effect in the DSF denaturation experiment, in which reduced WT-FBPase was stabilized by +12°C to a T_m_=54.68 ±0.68°C (Table 1). Conversely, DTT treatment of PROSS-FBPase caused a marginal destabilization with -5°C to -9°C drop in thermal stability to a T_m_=75.51 ±0.3°C. We conclude that PROSS redesign of sequence increased the intrinsic stability of the enzyme *per se*, maintained its affinity to substrate and cofactors, and favored a conformation that is intrinsically more active and less dependent on redox external activation that WT-FBPase. The beneficial or adverse effects of reducing agent on natural or redesigned enzyme owns both to a multiplicity of redox controls, to the increase in PROSS-FBPase stability in heterologous host, and to the modification of cysteines content by PROSS (*vide infra*).

Besides, the higher intrinsic activity of redesigned FBPase is consistent with the apparent molecular weight of the purified enzymes (Figure 2B), namely the homo-dimeric elution profile of WT-FBPase (apparent Mw=70 kDa, protomer Mw=40 kDa) and principally homo-tetrameric elution profile of PROSS-FBPase (apparent Mw=158 kDa). Most of reference literature reports FBPase to be active in the homo-tetrameric state, with a notable mechanistic demonstration of tetrameric cooperativity on the chloroplast isoform (Giudici-Orticoni et al., 1990). PROSS redesign of sequence had an apparent positive effect on the stabilization on homo-tetrameric quaternary structure, hence contributing to a higher intrinsic activity of purified recombinant FBPase *in vitro*.

Besides, we assayed the response of FBPase to varying pH, considering that the stroma alkalinization/acidification as a consequence of photosystems activity is correlated with a slow pH-induced conformational transition of chloroplastic FBPase (Gontero et al., 1984). In our hands, both WT- and PROSS-FBPase were indeed sensitive to pH changes in the 7-8 physiologically relevant window (Figure 3C) and with similar sensitivity profiles. In this instance, PROSS redesign did not induce a modification of properties compared to physiological reference.

### Crystal structure and computational modeling of PROSS-FBPase

In order to consolidate structural modeling and to facilitate future engineering of FBPase, we determined the experimental structure of PROSS-FBPase by X-ray crystallography. While we never succeeded thus far in growing crystals of WT-FBPase despite extensive screening of crystallization conditions, PROSS-FBPase did crystallize in several conditions of our standard, identical pipeline.

We tested diffraction on a series of crystals and collected complete diffraction datasets of PROSS-FBPase crystals in space group P 4_1_2_1_2, the best being at a resolution of 2.29 Å (Figure 4). We could determine the phases by molecular replacement with an Alphafold3 (AF3) model of the protomer and 2 copies of it in the asymmetric unit. AF3 model and the refined experimental structure align with an overall RMSD=0.459 Å, agreeing on the core secondary structures and the chain topology. The main regions of divergence are located in 53-NLTGVAGNQNVQGED-67 and 141-EE<u>C</u>PLDDMDDPQKMEQQ<u>C</u>I-159 that AF3 predicts to be ordered as a loop followed by a 12-residue long alpha helix, while the crystal structure is not defined in this region that is probably of high local disorder. Considering that region 141-153 contains candidate redox-regulatory disulfide-bridged cysteines identified in *Pisum sativum* as negative regulators stabilizing a local α-conformation (underlined), and taking into account the intrinsically active state of purified PROSS-FBPase, we consider that the crystal structure represents the catalytically active conformation trapped thanks to PROSS redesign strategy. Observed local disorder in the 141-159 segment so appears as a signature state of active FBPase.

**Figure 4.**
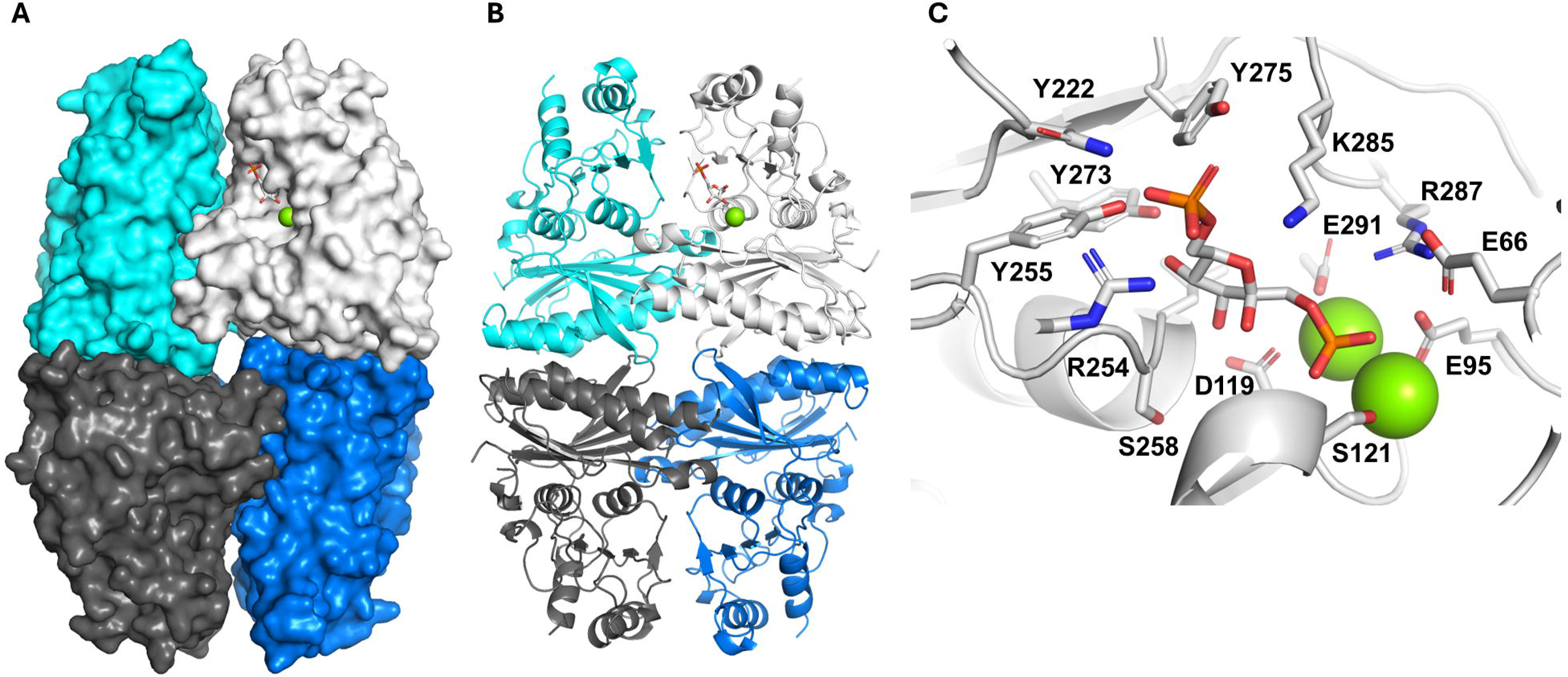
Crystal structure of PROSS fructose bisphosphatase, reconstituted in homo-tetrameric state from the dimeric asymmetric unit and the nearest neighbor dimer in crystal packing. A. Surface representation of PROSS-FBPase crystallographic model. Magnesium ions (green spheres) and fructose-1,6-bisphosphate (FBP, sticks) were placed by structural alignment with 1FBP crystal structure of full active site (RMSD=0.805 Å), as a guide to locate catalytic pockets. B. Similar to A with main chain cartoon representation of PROSS-FBPase crystallographic model. C. Catalytic pocket predicted by Boltz-2 with residues in polar bonding distance to FBP and Magnesium ions.

### Reconstitution of a model tetrameric FBPase bound to substrate

After refinement of the crystal structure and the reconstitution of its homo-tetrameric biological assembly with crystalline neighbors, we built computed a model of the protein bound to Magnesium cofactor and the FBP substrate (Figure 4A-B). Crystallographic dimer of PROSS-FBPase structurally aligns to the crystal structure 1SPI of *Spinacia oleracea* FBPase with RMSD=1.771 Å. Main differences arise from the relative orientations of subunits towards each other, with PROSS-FBPase dimer being opened with a wider angle than its spinach homolog. AF3 model, crystal structure 1SPI, and PROSS-FBPase crystal structure all agree on the main chain topology and quaternary organization, supporting a strong conservation of structure-function relationship among all photosynthetic FBPases. We thus computed with Boltz-2 a catalytic complex of PROSS-FBPase with Magnesium ions and FBP (Passaro et al., 2025) (Figure 4C). 16 residues contribute to the formation of the catalytic pocket: E66, E95, D116, D119, G120, S121, N222, R254, Y255, G257, S258, L259, Y273, Y275, K285, R287, E29. These residues should be primary targets for a future focused redesign of the enzyme catalytic pocket.

## Discussion

*Physicochemical gain-of-functions in FBPase, a core chloroplast metabolic enzyme* Starting from a poorly behaving recombinant WT-FBPase from *Chlamydomonas reinhardtii*, we used the redesign tool PROSS for extensive stabilization mutagenesis. We selected design8 from the standard automated computational pipeline, ordered synthesis and cloning of the corresponding gene for overexpression in bacteria, and prepared the corresponding PROSS-FBPase variant. Purified protein expressed highly in soluble state, and visibly remained soluble over several weeks at room temperature. Differential scanning fluorimetry (DSF) indicated that compared to thermolabile WT-FBPase, our PROSS-FBPase variant gained +40°C in thermal stability (T_m_>80°C), consistent with the prediction of an enthalpic stabilization. Dual denaturation temperature of PROSS-FBPase in the absence of added ligand, compared to the single denaturation temperature in the presence of substrate (FBP) or cofactor (Mg^2+^) indicates that purified *apo* protein is a mixture of 2 populations of different conformational states, that the addition of substrate or cofactor biases towards a single over-stabilized state.

*In vitro*, PROSS-FBPase is active in its specific reporter assay quantifying FBP hydrolysis function. The redesigned enzyme is purified in active form, that correlates with a homo-tetrameric population on SEC, and displays Michaelis-Menten kinetic parameters (K_M_=0.230 mmol/L and k_cat_=0.1 sec^-1^) similar to those of chloroplastic FBPase from the microalga Euglena. Recombinant PROSS-FBPase is not activated by chemical reducing agent, in striking contrast to recombinant WT-FBPase that purifies in an oxidized, inactive state. Redesign of the sequence apparently selected a sequence favoring the active form of the enzyme, probably through bias towards consensus sequences during the multiple sequence alignment of PROSS. Nevertheless, redesign FBPase has a similar activity in the same order of magnitude to the WT reference post-reduction, and also kept a natural sensitivity to pH.

### Insights into redox regulatory properties of chloroplastic FBPase from microalgae

FBPase from Chlamydomonas natively contains 9 cysteine residues, and is documented as a redox-responsive enzyme. Thioredoxin-dependent reduction of disulfide bridges in photosynthetic FBPase allows for a conditional activation of the CBBC when photosystems are active. Among the 9 cysteines of CrFBPase a multiplicity of redox regulation including nitrosylation, glutathionylation, and dithiol/disulfide exchanges were reported, with an open question regarding the combinatorial effects of simultaneous or sequential regulations. Interestingly, Chlamydomonas displays 2 additional cysteines compared to land plants like *Pisum sativum*, opening a question about the relative importance of these putative regulatory sites. Given the proximity of these 2 microalgae-specific cysteines to plant-generic cysteines in pentapeptide motifs 77-CLAS<u>C</u>-81 and 171-<u>C</u>AGYC-175 (Figure 1A), we propose that these sites can form local intramolecular disulfide bridges that contribute to FBPase regulation in *Chlamydomonas reinhardtii*.

PROSS redesign of CrFBPase suppressed these 2 microalgae-specific cysteines, respectively susbstituting them with serine (position 81, C81S) and alanine (position 171, C171A), returning the redesign sequence to a consensual non-microalgal, plant isotype (Figure 1A). These positions correspond to non-conserved cysteine sites that are specific to microalgae and absent in land plants and resulting PROSS-FBPase actually behaves strikingly differently than WT-FBPase in recombinant state: absence of apparent precipitation at production stages, long-term activity, and a dominant homo-tetrameric state at the final purification step (Figure 2B). Accordingly, PROSS-FBPase purifies as an active protein without additional treatment, in contrast to WT-FBPase that requires chemical reduction to reach a catalytically active state (Figure 3B). This distinct behavior leads us to conclude that the microalgae-specific C81 and C171 cause intrinsic instability correlated with low activity in the recombinant WT-FBPase, that can only be compensated by disulfide reduction after purification. PROSS redesign attributes a negative regulatory role to microalgae-specific cysteines, and provides a tool to evaluate the function of the remaining plant-generic redox control in the microalgae background. Besides, the stabilization of the homo-oligomeric state into active tetramer in PROSS variant in the absence of redox modification is reminiscent of the redox control of dimer/tetramer activating transition of CrSBPase (Le Moigne et al., 2025). It is likely that C86 and C171 are dynamically redox modified, thus switching inactive dimers to active tetramers, with a catalytically-beneficial local disorder conditioned by the reduction of disulfides. In support of the hypothesis of a disulfide control over tetramerization of CrFBPase, residue 86 is exposed at the dimer-to-dimer interface in PROSS-FBPase crystal structure. The redox state of the 2 microalgal-type cysteines should be probed further in the WT background to demonstrate their implication in this new layer of CBBC regulation.

### Contribution to the engineering of photosynthetic carbon fixation

The gain in thermal stability through PROSS computational redesign can solve an intrinsic dependence of the WT-protein over the chaperone system for high temperature buffering. Indeed, *Chlamydomonas reinhardtii* being a unicellular microalga living in natural small bodies of water are exposed to environmental variations of temperature that can easily exceed T_m_ of WT-FBPase, especially in the context of global warming. Substitution of thermostabilized variant could thus relieve the cell from a dependence over heat-shock chaperones, saving energy or proteostasis potential for other functions.

It was recently reported that AI-redesigned protein sequences are better starting points in directed molecular evolution (Krasnow et al., 2026). We describe here a relevant entry point for the re-evolution of FBPase. As a first attempt at improving PROSS-FBPase, we submitted it to FuncLib for targeted refinement of its catalytic pocket residues (Khersonsky et al., 2018),(Lipsh-Sokolik and Fleishman, 2025). Conceptually parallel to PROSS algorithm, FuncLib consists in a massive multiple alignment search of multipoint amino acid variants for improved predicted stability (free-energy) of the redesigned protein. FuncLib-PROSS-FBPase variant behaved similarly to PROSS-FBPase for expression and purification. However, it unexpectedly appeared devoid of any enzymatic activity in our *in vitro* reporter assay. Besides, FuncLib-PROSS-FBPase differential scanning fluorimetry profile remained unchanged in the presence or absence of substrate, which would be explained if the protein was locked into a highly stabilized Michaelis complex, with FBP and Magnesium irreversibly taken from the bacterial expression host. Then, the paradoxical loss of functional improvement when the catalytic pocket is redesigned at its highest stability would illustrate the Sabatier principle, *i.e.* that a catalyzer should bind the intermediate state of reaction with an affinity that still allows for conformational rearrangements during catalysis and the final release of products.

### Limitations of the study

This work is primarily focused on one alternative variant to the natural version of FBPase. The computational strategy, PROSS, was applied in parameters of reference, with the single solution pross8 taken for experimental validation. An interesting complement to the reported results would be a systematic testing of proteins from design1 to design9, incrementally adding modifications to the reference sequence up to the point when the gain-of-function is considered significant. Alternatively, another validation strategy would be to revert mutations one-by-one from design8, until the gain-of-functions are lost and the protein behaves as the WT. This limit is overall linked with the weak rational that we recognize on the mutation-function relationship, because the computational redesign of PROSS was systematic and extensive by nature.

Regarding the biological potential of the modified sequence, one may expect that PROSS-FBPase would only partly fit within a chloroplast proteome, mainly because of its gained independence towards redox activation. Resulting metabolism would be a permanent drive to anabolism than is likely detrimental in natural conditions. Future work on *e.g.* Chlamydomonas transformed with PROSS-FBPase expression transgenes should be tested in growth conditions like continuous light in photobioreactor or in the presence of additional carbon source (*e.g.* TAP medium). The intrinsic resistance of PROSS-FBPase to thermal denaturation *in vitro* would confer a resistance to natural turnover, potentially unbalancing proteostasis. Finally, it is possible that an over-stabilized enzyme would lose its evolutionary potential in natural genomic background, reducing its interest for *in vivo* selection of beneficial traits.

## Conclusions

This study reports an extensive redesign of the sequence of chloroplastic-type FBPase with computational-guided mutagenesis. Through comparison of stability and activity with the natural (WT) sequence, this study describes an unexpected multiplicity of redox controls over this enzyme that fits at the metabolic crossroads between photosynthetic carbon fixation and the general metabolism through neoglucogenesis and sucrose biosynthesis.

The additional layers of redox-posttranslational modifications exerted over FBPase from *Chlamydomonas reinhardtii* green microalga parallels previous reports on another CBBC enzyme from the same organism, phosphoglycerate kinase (PGK) (Michelet et al., 2013),(Morisse et al., 2014). The unique microalgae redox-dependence in both FBPase and PGK indicates a tighter control on both the reductive and the regenerative phases of the CBBC in eukaryotic microalgae, plausibly linked with their adaptation to changing environment as single cell organisms. This multiplicity of controls should also be interpreted with the variety of redox post-translational modifications (Pérez-Pérez et al., 2017),(Zaffagnini et al., 2019) and other types of modifications (Wang et al., 2014),(McConnell et al., 2018),(Serre et al., 2020),(Füßl et al., 2022) to characterize how the dynamic modification of chloroplast proteome exerts a tight control of carbon budget in the whole green lineage.

The reported crystal structure for a new CBBC-related enzyme contributes to the achievement of the molecular description at high resolution of this important pathway, unified within the model microalga *Chlamydomonas reinhardtii* taken as a reference for *Viridiplantae* (Taylor et al., 2001),(Pasquini et al., 2017),(Gurrieri et al., 2019),(Le Moigne et al., 2020),(Mattioli et al., 2022),(Le Moigne et al., 2022),(Meloni et al., 2024),(Le Moigne et al., 2025),(Meloni et al., 2026). Integration of all structural models at high resolution, including the photosynthesis-specific modes of redox-, pH-, and divalent metal-dependent regulations, along with the oligomerization switches driving homotropic cooperativity, supports a global modeling of the CBBC as a system.

## Declarations

### Availability of data and materials

Data and materials are available upon request to corresponding author (JH).

### Fundings

Institut Universitaire de France (JH)

Agence Nationale de la Recherche ANR-24-CE45-7686 DEFINE

Agence Nationale de la Recherche ANR-25-CE11-2985 CALVINASSEMBLY Partenariat Hubert Curien Galileo 52406/54359SA

Institut de l’Océan de l’Alliance Sorbonne Université (CQP)

### Authors’ contributions

CQP, TC, and JH designed and conducted experiments, including the preparation of recombinant proteins, the characterization of their *in vitro* parameters, their crystallization and the collection of X-ray diffraction datasets. CQP, TC, and JH analyzed resulting data. JH conceptualized the study, acquired fundings, designed methodology, solved the crystallographic data, and supervised the study. All authors contributed to the preparation, writing and revision of the manuscript. All authors read and approved the final version of the manuscript. In accordance with BMC guidelines, large language models (LLMs) were not used for the writing of the manuscript.

## Acknowledgements

We acknowledge Pr. Sarel Fleishman (Laboratory of Protein Design, Weizmann Institute of Science, Rehovot Israel) for providing privileged access to PROSS and FuncLib softwares online. We thank the Institut de Biologie Physico-Chimique (CNRS FR 550) for access to crystallization facilities, SOLEIL for provision of synchrotron radiation facilities at beamlines Proxima-1 and Proxima-2A, and local contact Dr. Pierre Montaville for help with data collection and treatment. We thank Pr. Mirko Zaffagnini, Pr. Paolo Trost, Pr. Francesca Sparla, Pr. Simona Fermani, Dr. Libero Gurrieri, and colleagues from the department of Pharmacology and Biotechnology, University of Bologna, for insightful discussions.

